## Supplemental text with figures for "Predicting scale-dependent chromatin polymer properties from systematic coarse-graining"

Sangram Kadam,[1,*] Kiran Kumari,[1] Vinoth M,[1] Shuvadip Dutta,[2] Mithun K. Mitra,[2] and Ranjith Padinhateeri[1,3,†]
[1]*Department of Biosciences and Bioengineering, Indian Institute of Technology Bombay, Mumbai 400076, India*
[2]*Department of Physics, Indian Institute of Technology Bombay, Mumbai 400076, India*
[3]*Sunita Sanghi Centre for Aging and Neurodegenerative Diseases, Indian Institute of Technology Bombay, Mumbai 400076, India*

#### S1. SUPPLEMENTARY METHODS

##### A. Model-I: Fine-grained chromatin model with 200bp resolution chromatin

We consider chromatin as a bead-spring chain polymer made of $N$ beads. Each bead is a 200bp chromatin—nucleosome + linker DNA—connected by springs of stiffness $k_s$. Let $\sigma$ be the diameter of the bead. Since each bead is essentially a nucleosome+linker DNA, we estimate $\sigma \approx 21$nm (see below). We used the publicly available Micro-C data from (Hsieh et al., 2020) for mouse embryonic stem cells as input. The Micro-C contact matrix is KR normalized (Hsieh et al., 2020) and further scaled by the sum of all the row elements of the matrix to get the contact probability values $P_{ij}$.

We follow a two-step process to generate an ensemble of configurations consistent with the Micro-C data. First, we define a set of prominent contacts from the Micro-C data. We define the prominent contacts matrix as follows.

$$P_{ij}^{pr} = \begin{cases} P_{ij} & \text{if } P_{ij} > \mu(|i-j|) + s.d.(|i-j|), \\ 0 & \text{otherwise} \end{cases} \tag{1}$$

Here $\mu(|i-j|)$ and $s.d.(|i-j|)$ are the mean and standard deviation of contact probability values $P_{ij}$ for bead pairs $i$ and $j$ at constant separation $|i-j|$ along the contour. These represent the $P_{ij}$ values lying parallel to the diagonal of the matrix. We chose to define prominent contacts by considering the mean and standard deviation of the same $|i-j|$ bead-pairs because for a homogeneous polymer, purely due to its polymer nature, the contact probability depends only on $|i-j|$.

Then we generate an ensemble of 1000 binary contact matrices $C$, which satisfy the contact probability $P^{pr}$. To achieve this, in each matrix, we set $C_{ij} = 1$ if $r_n < P_{ij}^{pr}$, else $C_{ij} = 0$, where $r_n$ is a uniform random number between 0 and 1. Here the matrix element $C_{ij} \in \{0, 1\}$ represents contact between $i^{th}$ and $j^{th}$ bead in a given configuration. Then we connect the bead pairs $i$ and $j$ having $C_{ij} = 1$ through harmonic springs, having energy

$$E_{ij}^{mc} = \frac{k_s}{2} \sum_{\substack{i,j \\ j>i+1}} C_{ij}(|\vec{r}_i - \vec{r}_j| - \sigma)^2. \tag{2}$$

We generated 1000 such configurations (corresponding to 1000 contact matrices $C$) and independently equilibrated them using Langevin Dynamics simulations with LAMMPS. The total energy of the polymer is given by

$$E = \frac{k_s}{2} \sum_{i=1}^{N-1} (|\vec{r}_i - \vec{r}_{i+1}| - \sigma)^2 + \sum_{\substack{i,j \\ j>i}} E_{WCA}(|r_{ij}|) + E_{ij}^{mc}. \tag{3}$$

---

*Electronic address:
†Electronic address:

The first term represents the polymer connectivity via springs, and the second term represents the steric repulsion given by

$$E_{WCA}(r) = \begin{cases} 4\epsilon\Big[\Big(\frac{\sigma}{r}\Big)^{12} - \Big(\frac{\sigma}{r}\Big)^{6}\Big] + \epsilon & r < 2^{1/6}\sigma, \\ 0 & r \geqslant 2^{1/6}\sigma. \end{cases} \tag{4}$$

As discussed above, the third term in Eq. 3 represents the intra-chromatin interaction between different segments based on the Micro-C data.

In the second step, we used the equilibrium configurations obtained from the above step. For each pair of beads $i$ and $j$, we inserted the spring contacts $C_{ij} = 1$ in the $P_{ij}$ fraction of configurations having the smallest 3D distance $r_{ij}$ between the pair. Then we equilibrated these configurations similar to the first step using this updated contact matrix $C$.

We begin by initializing the polymer configuration with a self-avoiding walk. We take a step-wise approach to ensure the polymer configuration is stable and does not experience large forces from extra contacts through Micro-C interactions between far-away beads. We introduce extra contacts gradually. First, we start with local contacts between bead pairs $i$ and $j$ that are close together (with a separation of $|i-j| < 5$) and equilibrate the polymer for $10^5$ steps. In subsequent steps, we introduce remaining contacts with larger separations ($|i-j| < 10$, $|i-j| < 25$, and so on), relaxing the polymer for $10^5$ steps after each step. Once all contacts have been added, we equilibrate the polymer for $6 \times 10^6$ time steps. During the last $3 \times 10^6$ steps, we sample the data every 50000 steps to analyze the behavior of the polymer in more detail.

We also simulated Self Avoiding Walk (SAW) polymer, ideal Gaussian chain, and polymer globule with attractive interactions for comparison with chromatin polymer. Unlike the fine-grained chromatin model, these polymers do not have extra spring contacts from the contact map. These polymer models are simulated for $N = 400$ beads using Langevin simulations in LAMMPS. The detailed energies are given below.

**SAW Polymer:** The polymer consists of $N$ beads connected by harmonic springs. The total energy is given by

$$E = \frac{k_s}{2}\sum_{i=1}^{N-1}(|\vec{r}_i - \vec{r}_{i+1}| - \sigma)^2 + \sum_{\substack{i,j \\ j>i}} E_{WCA}(|\vec{r}_{ij}|). \tag{5}$$

Here $E_{WCA}$ is used for excluded volume effect given by Eq. 4. Here the value of spring constant $k_s = 10$, the size of the bead $\sigma = 1$, and the energy parameter value in $E_{WCA}$ is $\epsilon = 1$.

**Ideal Gaussian chain:** The ideal Gaussian chain polymer has $N$ beads connected by harmonic springs without excluded volume interaction. In other words, it is the same as SAW with $E_{WCA} = 0$. The value of the spring constant is $k_s = 10$.

**Globule:** We simulate polymer globule using a bead-spring polymer with attractive LJ interaction given by

$$E_{LJ}(r) = \begin{cases} 4\epsilon\Big[\Big(\frac{\sigma}{r}\Big)^{12} - \Big(\frac{\sigma}{r}\Big)^{6}\Big] & r < 2.5\sigma, \\ 0 & r \geqslant 2.5\sigma. \end{cases} \tag{6}$$

The total energy is given by

$$E = \frac{k_s}{2}\sum_{i=1}^{N-1}(|\vec{r}_i - \vec{r}_{i+1}| - \sigma)^2 + \sum_{\substack{i,j \\ j>i}} E_{LJ}(|\vec{r}_{ij}|). \tag{7}$$

Here the spring constant is $k_s = 10$, bead size $\sigma = 1$, and the LJ energy parameter $\epsilon = 1$.

#### B. Model-II: Model with explicit linker DNA

This is a bead-spring model with explicit linker and nucleosome details (Fig.S2(b)). In this model, the chromatin polymer has two types of beads – linker DNA bead ($D$) and nucleosome bead ($N_u$). Each DNA bead represents

$\approx$ 10bp of linker DNA and has a size $\sigma_d = 3.4nm$, while the nucleosome beads are of size $\sigma_n = 3\sigma_d = 11.2nm$. Neighboring nucleosome beads are connected by five linker DNA beads (see Fig.S2(b)). The beads in this polymer are connected by a harmonic spring, whose potential is given by,

$$E_s = \frac{k_s}{2}(|\vec{r_i} - \vec{r}_{i+1}| - r_0)^2, \tag{8}$$

here $\vec{r_i}$ and $\vec{r_{i+1}}$ are position vectors of neighboring beads, $r_0$ is the equilibrium bond length, and $k_s$ is the spring constant. The equilibrium bond lengths for $D - D$ and $D - N_u$ bonds are $1\sigma$ and $2\sigma$, respectively. The value of $k_s$ is taken as $10K_BT/\sigma_d^2$. Since we have explicit linker DNA and nucleosomes with entry-exit angles in this model, the bending rigidity is modeled by the potential,

$$E_b = \frac{k_b}{2}(1 - cos(\theta_i - \theta_0)), \tag{9}$$

where $\theta_i$ is the angle between $i^{th}$ and $(i+1)^{th}$ bond vectors , $\theta_0$ is the equilibrium bond angle and $k_b$ is the bending stiffness of the angle. In our simulations, the $k_b$ for $D - D - D$ and $D - N_u - D$ angles are taken as $28k_{\rm B}T$ and $3.8k_{\rm B}T$, respectively. The bending rigidity of the linker DNA beads is taken considering that it is highly rigid, whereas $D - N_u - D$ bending stiffness is taken in accordance with the experimental data(Beel et al., 2021). We assumed a bending stiffness for $D - N_u - D$ ($k_b = 3.8k_{\rm B}T$) such that angle distributions around $60°$ is similar to (Beel et al., 2021). The equilibrium angle for $D - D - D$ is 180° while for $D - N_u - D$, the angle is 60° and 90°. All other non-bonded beads interact with WCA potential (Eq.4). Then, we performed classical Langevin dynamics simulations for this model in LAMMPS molecular dynamics package.

We computed the average $R_g$ of 1kb chromatin polymer from this model and compared it with model-I (Fig. S2(c)). The average $R_g$ of 1kb from our model-I is comparable to what we get from detailed model-II. This suggests that the choice of NL bead in model-I is sensible. In real chromatin, there exists linker length variability. To test whether this variability could affect the compaction, we used the number of linker beads from a Gaussian distribution with a mean of five linker DNA beads (50bp) and the standard deviation of 1 linker DNA bead (10bp). The average $R_g$ of the 1kb chromatin polymer with linker length variability is still comparable to our model-I (Fig. S2(c,d)). Moreover, the mean linker length in euchromatic regions can be higher than that of the heterochromatic regions. To understand how this difference in mean linker length would affect the size of 1kb chromatin, we varied the mean to four (40bp) and six (60 bp) linker DNA beads, keeping the standard deviation the same as before (1 linker DNA bead, 10bp). The change in size (mean $R_g$) due to this difference in mean linker length is reported in Fig S2(d).

#### C. Iterative Boltzmann inversion method

We estimate the nonbonded interactions for the Arsg locus at different coarse-graining levels. The CG polymer consists of $N/n_b$ number of beads; the spring coefficient values ($K_{cg}$) are taken from Fig. 3(h) in the main manuscript. We implement the iterative Boltzmann inversion method as follows (see Fig. S12). The distribution of the distance between nonbonded beads computed from the original fine-grained model for the corresponding level of coarse-graining is taken as the target distribution ($P_{\rm target}(r)$). We use zero interaction ($V_{i=0}^{nb} = 0$) as the starting guess for the potential. We run the Metropolis Monte Carlo simulations for each iteration with the following energy.

$$E = \frac{K_{cg}}{2}(|\vec{r_i} - \vec{r}_{i+1}| - l_{cg})^2 + \sum_{\substack{i,j \\ j>i+1}} V^{nb}(|\vec{r}_{ij}|). \tag{10}$$

We equilibrate the polymer and then compute the distribution of the distance between nonbonded beads. We check for convergence by computing the Kullback-Leibler Divergence given by

$$D_{KL}(P_{\rm target} \mid\mid P_i) = \sum_r P_{\rm target}(r)\ ln\left(\frac{P_{\rm target}(r)}{P_i(r)}\right)$$

If the distribution has not converged, we update the potential for the next iteration as follows.

$$V_{i+1}^{nb}(r) = V_i^{nb}(r) + \alpha(r)\ k_BT\ ln\left(\frac{P_i(r)}{P_{\rm target}(r)}\right) \tag{11}$$

Here $\alpha(r) = 0.2\ e^{-r^2/2}$ is a decaying function to ensure that the resulting potential is short-range. Fig. S13(a) shows the Kullback-Leibler Divergence as a function of iteration, which converges to zero.

We also use a functional form to fit the obtained potential from the above IBI method. The functional form used is given by

$$V_{\text{soft}}(r) = \begin{cases} V_0\big[1 - \big(\frac{r}{r_m}\big)^{\eta_1}\big]^{\eta_2} - \epsilon & r < r_m, \\ \frac{1}{2}\epsilon\big[\cos(\mu r^2 + \nu) - 1\big] & r_m \leqslant r < r_c, \\ 0 & r \geqslant r_c. \end{cases} \tag{12}$$

The first part of the equation ($r < r_m$) represents the repulsive part of the potential (Fujishiro and Sasai, 2022), while the second part ($r_m \leqslant r < r_c$) shows the attractive part (Soddemann et al., 2001). The values of $\mu$ and $\nu$ are set by solving the following two equations.

$$\mu\ r_m^2 + \nu = \pi$$

$$\mu\ r_c^2 + \nu = 2\pi$$

These equations ensure that the value of the potential is $-\epsilon$ at $r = r_m$ and zero at $r = r_c$, ensuring continuity and differentiability at these points. The interaction energy is zero for the long-range interactions ($r > r_c$). Values of all the fitting parameters used for different values of $n_b$ are reported in Table S2.

#### D. Quantities measured

- **Contact probability:** The contact probability between any bead-pair $i$ and $j$ is computed as

$$P_{ij} = \frac{1}{N_{conf}} \sum_{k=1}^{N_{conf}} H(r_{cut} - r_{ij}).$$

Here, $N_{conf}$ is the total number of polymer configurations, $H(r)$ is the Heaviside step function, and $r_{cut}$ is the cut-off distance below which two beads are considered to be in contact. We use $r_{cut} = 1.2\sigma$ as the cut-off for defining contact in the fine-grained model.

- **Stratum-adjusted correlation coefficient (SCC):** The contact probability between any two genomic regions is correlated to the genomic separation between them. Standard methods like Pearson correlation give a higher correlation between different contact maps due to these biases, which can be misleading. Hence we use a stratum-adjusted correlation metric (Yang et al., 2017) that stratifies the data based on the genomic distance eliminating the distance dependence, to compute the correlation between experimental and simulation contact maps. Before computing the SCC, we use 2D mean filter smoothing to smooth the contact map, where each contact probability value $P_{ij}$ is replaced by the mean of contact probabilities in the neighborhood (a square of size $2h + 1$ centered around $P_{ij}$ in the contact matrix). We use $h = 5n_b(1kb)$ for the smoothing algorithm. Then we divide the data into $K$ different strata based on the distance along the contour $(|i - j|)$. $K = 100$ is the number of strata. For each stratum, we compute the Pearson correlation coefficients between the $\log(P_{ij})$ values from the experiment and simulation. The weighted sum of these coefficients gives the stratum-adjusted correlation coefficient (SCC)(Yang et al., 2017). As a control, we measure the SCC between the SAW polymer and experimental contact map of the Ppm1g locus, which is $\approx 0$, suggesting that the SCC gives an accurate measure accounting for the distance dependence. The SCC values for all chromatin loci are given in Table S1.

We simulate the fine-grained model and generate an ensemble of equilibrium configurations. For each of these configurations, we coarse-grain the system by considering $n_b$ fine-grained beads as one coarse-grained (CG) bead (Main text Fig. 2(a)). We measure the following quantities for the CG polymer.

- **Radius of gyration:** The radius of gyration $R_g$ is used to quantify the size of polymer segments (as a function of the number of beads $n_b$ used for coarse-graining). The average radius of gyration of the polymer segment of $n_b$ NL beads forming $i^{th}$ CG bead is given by

$$R_g^i = \sqrt{\frac{1}{n_b} \Big\langle \sum_{k=i}^{i+n_b-1} |\mathbf{r}_k - \mathbf{r}_{com}^i|^2 \Big\rangle}.$$

Here angular brackets represent the ensemble average, $\mathbf{r}_k$ is the position vector of $k^{th}$ NL bead, and $\mathbf{r}_{com}^i$ is the position vector of center of mass of the $i^{th}$ CG bead given by,

$$\mathbf{r}_{com}^i = \frac{1}{n_b} \sum_{k=i}^{i+n_b-1} \mathbf{r}_k.$$

We also compute the total average radius of gyration $\langle R_g \rangle$, which is averaged over different genomic locations for a given coarse-graining $(n_b)$, given by

$$\langle R_g \rangle = \frac{1}{N - n_b + 1} \sum_{i=1}^{N-n_b+1} R_g^i.$$

- **Bond length between coarse-grained beads:** We have calculated the equilibrium bond length for adjacent coarse-grained beads (see Main text Fig. 2(a)) for various coarse-graining sizes $(n_b)$. The equilibrium bond length between $i^{th}$ and $(i+1)^{th}$ CG beads is calculated as

$$l_{cg}^i = \big\langle \mid \mathbf{r}_{com}^i - \mathbf{r}_{com}^{i+1} \mid \big\rangle.$$

Here $\mathbf{r}_{com}^i$ and $\mathbf{r}_{com}^{i+1}$ are the position vectors of $i^{th}$ and $(i+1)^{th}$ CG beads respectively.

We also compute the total average bond length $\langle l_{cg} \rangle$ for a given coarse-graining level $(n_b)$ by averaging over the genomic locations.

- **Spring constant:** We assume the consecutive coarse-grained beads to be connected by harmonic springs. The corresponding spring constant for the corresponding coarse-graining level ($n_b$) is calculated as

$$K_{cg} = \frac{k_B T}{\langle l_{cg}^2 \rangle - \langle l_{cg} \rangle^2}.$$

- **Bond angle between the coarse-grained beads:** The equilibrium bond angle between consecutive three coarse-grained beads ($i^{th}$, $(i+1)^{th}$ and $(i+2)^{th}$) is calculated using

$$\theta_{cg}^i = \langle cos^{-1}(\hat{l}_{cg}^i \cdot \hat{l}_{cg}^{i+1}) \rangle.$$

  Here $\hat{l}_{cg}^i$ and $\hat{l}_{cg}^{i+1}$ are the bond vectors (between consecutive CG beads) of unit magnitude given by

$$\hat{l}_{cg}^i = \frac{\mathbf{r}_{com}^{i+1} - \mathbf{r}_{com}^i}{\mid \mathbf{r}_{com}^{i+1} - \mathbf{r}_{com}^i \mid}.$$

- **Dihedral angle between the coarse-grained beads:** For four consecutive CG beads ($i$, $i+1$, $i+2$, and $i+3$) along the polymer, the dihedral angle $\phi_{cg}^i$ between them is the angle between two intersecting planes – plane consisting first three beads ($i$, $i+1$, and $i+2$) and plane consisting of last three beads ($i+1$, $i+2$, and $i+3$). We used the VMD software package to compute the dihedral angles(Humphrey et al., 1996).

**Parameters:** The Brownian dynamics simulations were performed using LAMMPS with LJ units based on energy ($k_B$T), mass ($m$), and distance $\sigma$. The spring constant for bonded beads was taken to be $k_s = 10$. The simulation was performed using a simulation box with shrink-wrap boundary conditions. The LAMMPS parameters were chosen as $damp = 1$, time step $\Delta t = 0.005$, and temperature $T = 1.0$.

### S2. SUPPLEMENTARY TABLES

| Locus Name | Chromosome | Start position (Mb) | End position (Mb) | SCC |
|---|---|---|---|---|
| Arsg | 11 | 109.42 | 109.67 | 0.90 |
| Alpha globin | 11 | 32.05 | 32.50 | 0.73 |
| Sox2 | 3 | 34.63 | 34.78 | 0.91 |
| Ppm1g | 5 | 31.17 | 31.25 | 0.95 |
| Nanog | 6 | 122.725 | 122.825 | 0.94 |
| Cbx8 | 11 | 119.0 | 119.1 | 0.98 |
| Gm29683 | 7 | 70.24 | 70.40 | 0.90 |
| Hoxa | 6 | 52.125 | 52.325 | 0.90 |
| Hoxb | 11 | 96.27 | 96.37 | 0.95 |
| Hoxc | 15 | 102.90 | 103.05 | 0.91 |

TABLE S1 The genomic locations of the various chromatin regions simulated in this study. The last column represents the stratum-adjusted correlation coefficient (SCC) values for each region.

| Coarse-graining size | $V_0$ | $\epsilon$ | $r_m$ | $\eta_1$ | $\eta_2$ | $r_c$ | $\mu$ | $\nu$ |
|---|---|---|---|---|---|---|---|---|
| $n_b = 5$ | 4.4 | 0.19 | 1.35 | 1.7 | 4 | 2.0 | 1.4 | 0.5 |
| $n_b = 10$ | 3.15 | 0.19 | 1.35 | 1.8 | 4.1 | 2.1 | 1.2 | 0.9 |
| $n_b = 25$ | 2.1 | 0.31 | 1.35 | 1.8 | 3 | 2.4 | 0.8 | 1.7 |
| $n_b = 50$ | 1.55 | 0.46 | 1.35 | 2.1 | 2.8 | 2.4 | 0.8 | 1.7 |

TABLE S2 Parameter values used in soft potential $V_{\text{soft}}$ for various levels of coarse-graining. The values of the parameters $\mu$ and $\nu$ are dependent on the values of $r_m$ and $r_c$.

### S3. SUPPLEMENTARY FIGURES

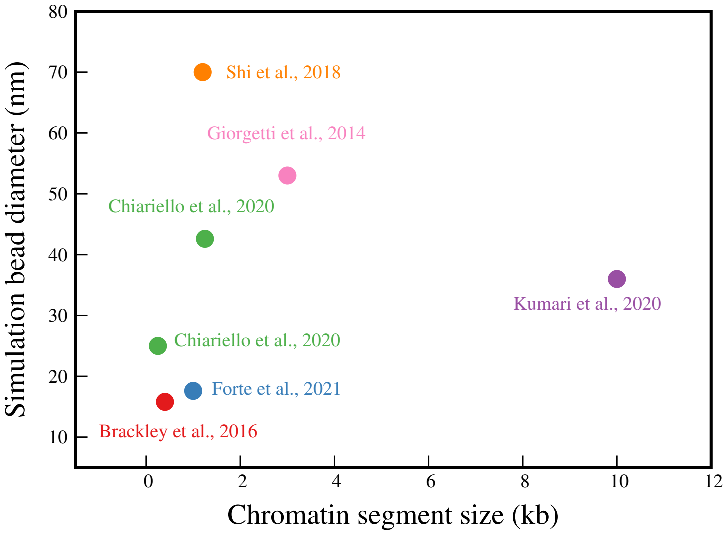


FIG. S1 Diameter (physical size) of simulation beads estimated for different chromatin segment lengths in various published papers (Brackley et al., 2016; Chiariello et al., 2020; Forte et al., 2021; Giorgetti et al., 2014; Kumari et al., 2020; Shi et al., 2018) Note that the points are highly scattered suggesting the need of a systematic study to understand size of chromatin segments. A brief description of the context of the data points is as follows: some of these studies investigate human chromosomes and estimate the same bead dimensions independent of the epigenetic state (Kumari et al., 2020; Shi et al., 2018). Shi et al. have studied Human Cell line GM12878 Chr 5 from 145.87 to 157.87 Mbps. Kumari et al. studied human chromosome 16 containing the alpha-globin gene and a few other genes such as LUC7L, K-562 (ON state) and GM12878 (OFF state). Forte et al. also estimated same bead-size for different epigenetic states studying a Pax6 locus in three mouse cell lines. Giorgetti et al. studied a mouse X chromosome region, presumably in an inactive epigenetic state. Brackley et al. studied mouse alpha globin locus. Chiariello et al. studied alpha and beta globin genes from mouse ES and erythroid cells.

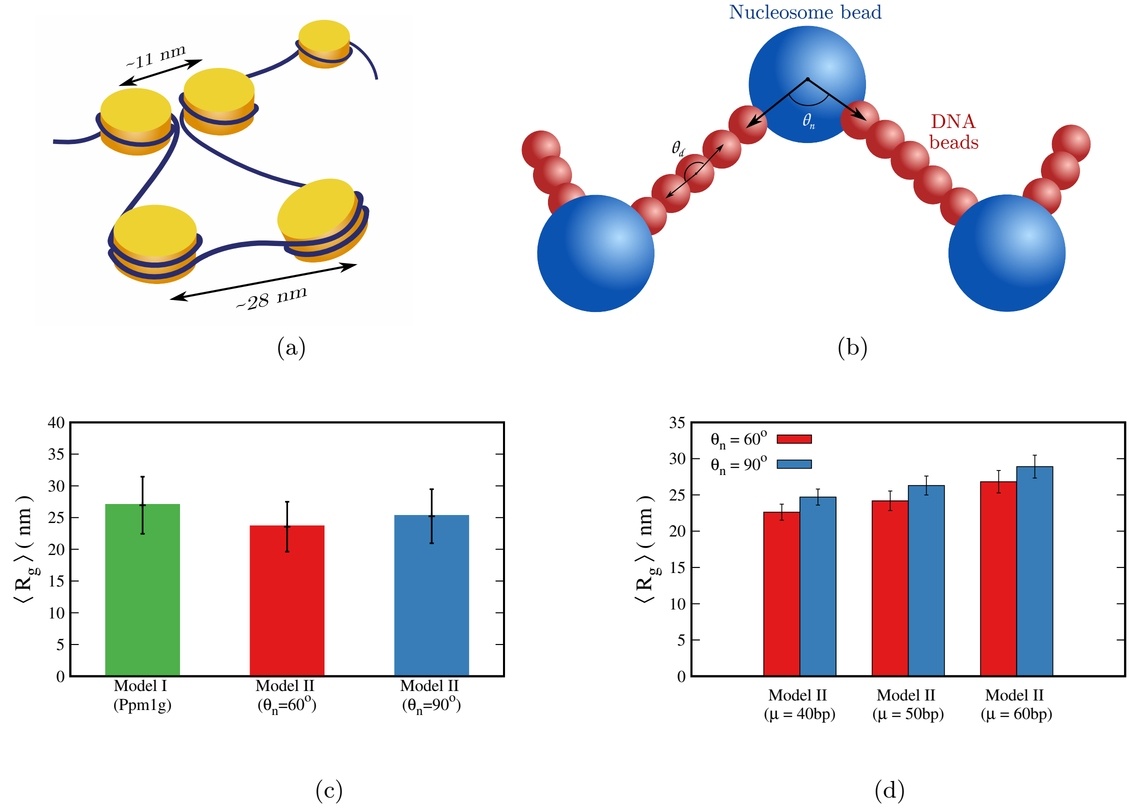


FIG. S2 (a) Geometric representation of separation between nucleosomes for two scenarios: when two far away nucleosomes touch each other (inter-bead distance 11nm) and nucleosomes with stiff linker DNA between them (inter-bead distance 28nm). (b)Schematic representing the model with nucleosomes and explicit linker DNA used in Model II (Sec. S1.B). (d) Correlation of contact probability from simulation and experiment for euchromatin and heterochromatin region. Representative snapshots of (c) Average radius of gyration $\langle R_g \rangle$ for 1kb chromatin computed from the model with NL beads (model-I), and compared it with a more detailed model (model-II) having explicit linker DNA and nucleosome-linker angles (simulated for two equilibrium nucleosome-linker angles: $\theta_n = 60^\circ$ and $\theta_n = 90^\circ$). Both Model I and Model II give comparable values of average radius of gyration for 1kb chromatin. (d) Radius of gyration of 1kb chromatin segment simulated by using Model II. The linker lengths in the simulation are taken from a Gaussian distribution with the specified mean ($\mu$ ) and a standard deviation of 10bp. Different mean values could mimic different chromatin states (heterochromatin, euchromatin). The standard deviation represents the variability in the number of nucleosomes. The results are presented for two nucleosome angles.

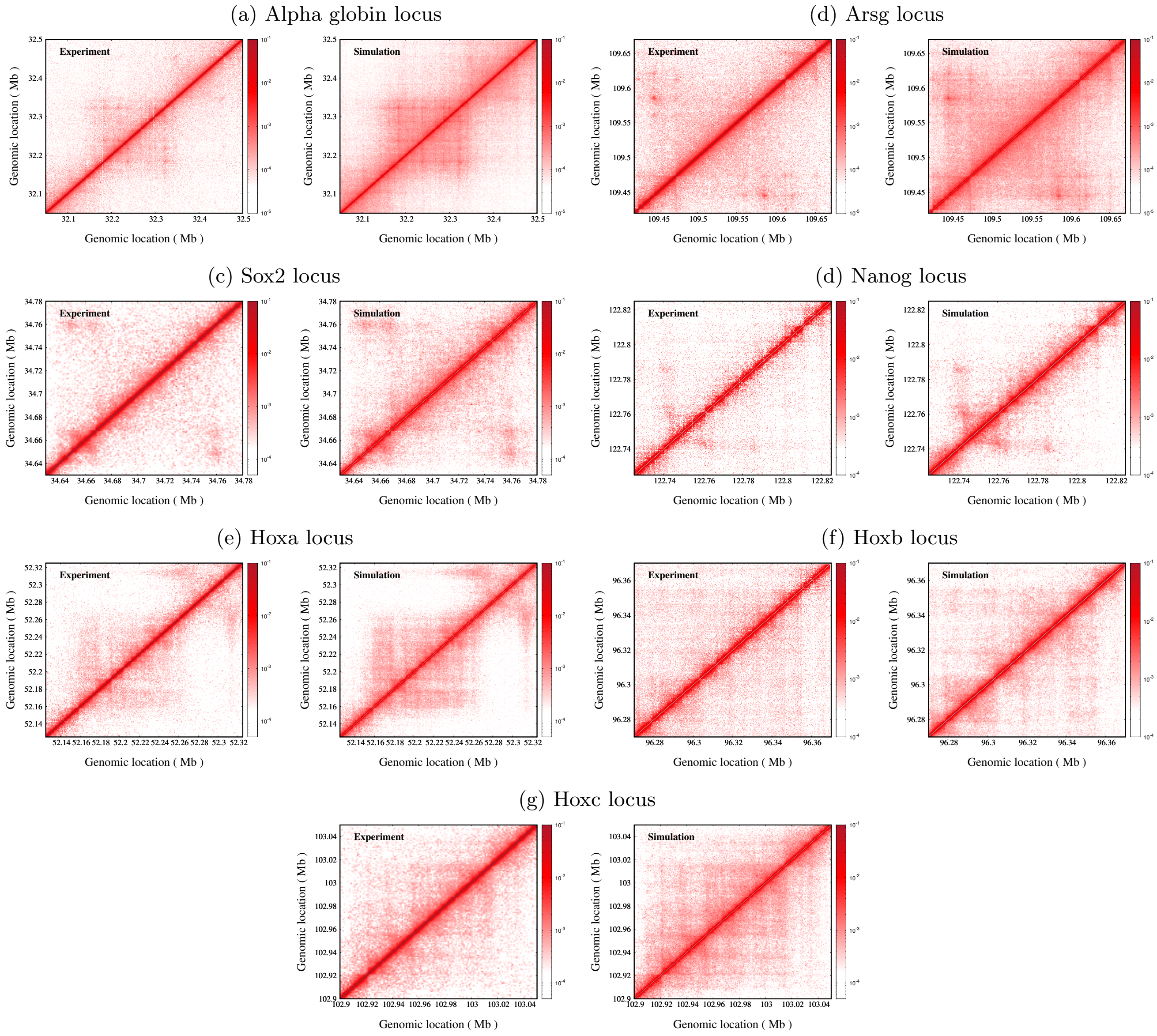


FIG. S3 Comparison of contact map from simulation with the Micro-C contact map for (a) alpha globin, (b) Arsg, (c) Sox2, (d) Nanog, (e) Hoxa, (f) Hoxb and (g) Hoxc loci.

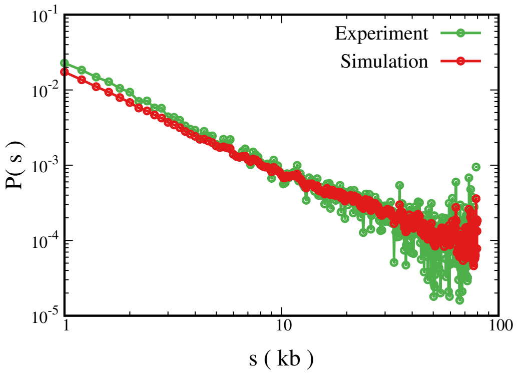


(a) Ppm1g locus

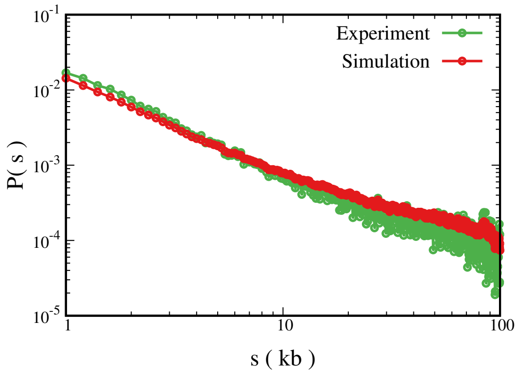


(b) Gm29683 locus

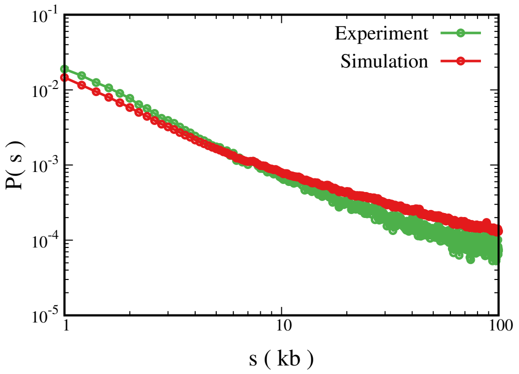


(c) Arsg locus

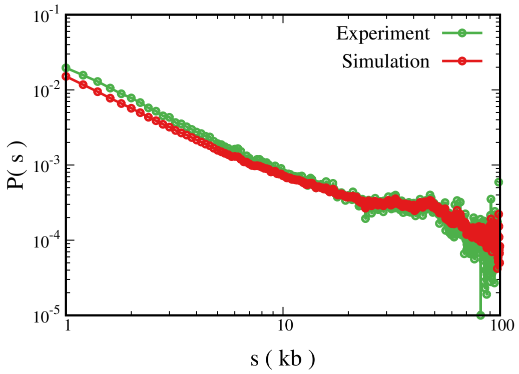


(d) Cbx8 locus

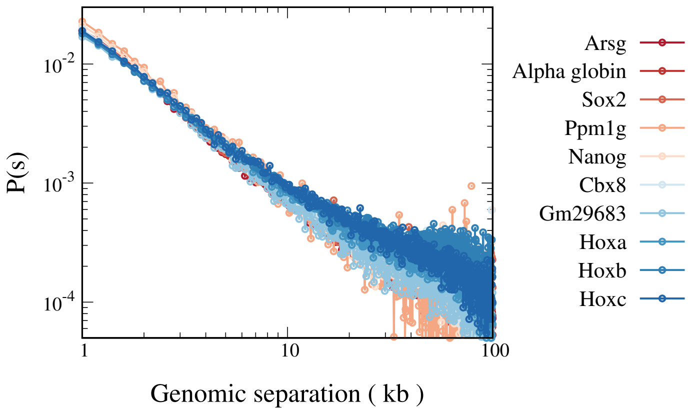


(e)

FIG. S4 Comparison of contact probability versus genomic distance from experiment and simulation for (a) Ppm1g, (b) Gm29683, (c) Arsg, and (d) Cbx8 loci. The stratum-adjusted correlation coefficients for all regions are reported in Table S1. (e) The experimental contact probability is plotted as a function of genomic distance for various euchromatic (shades of red) and heterochromatic (shades of blue) gene loci.

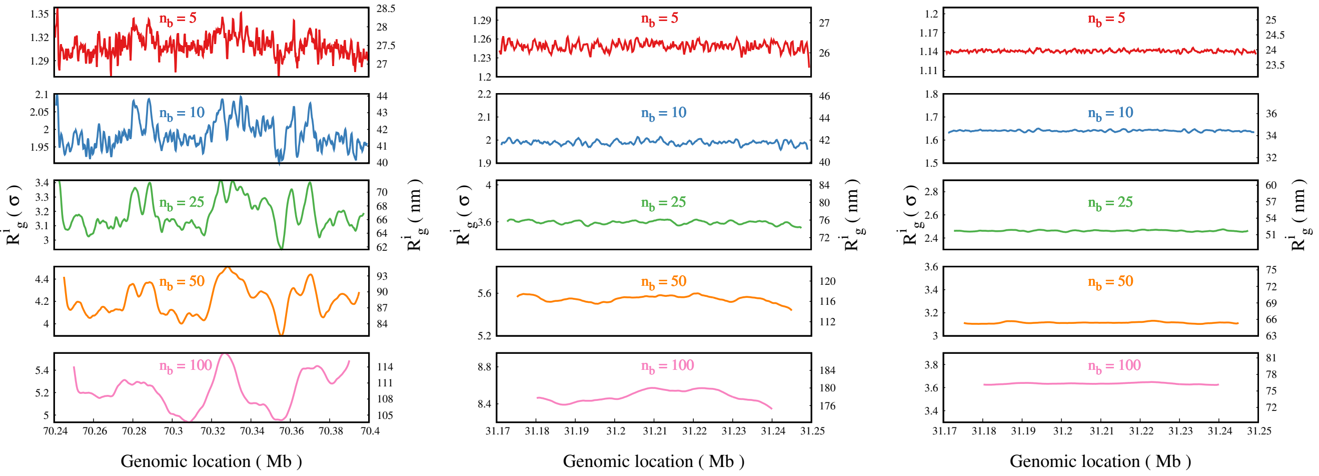


(a) Gm29683 locus (b) SAW (c) Globule

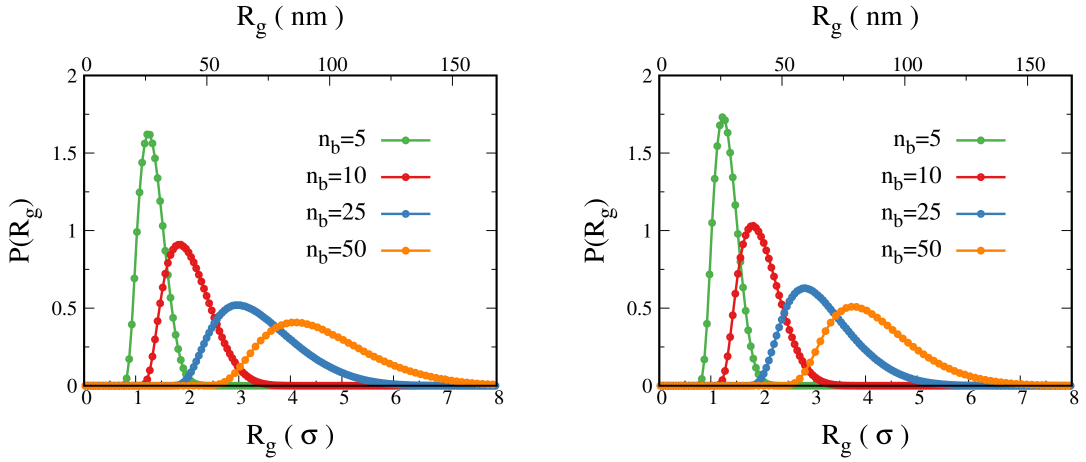


(d) Arsg locus (e) Gm29683 locus

FIG. S5 Average radius of gyration of CG beads as a function of genomic location for (a) Gm29683 locus, (b) SAW and (c) globular polymer. Distribution of $R_g$ for (d) Arsg locus and (e) Gm29683 locus.

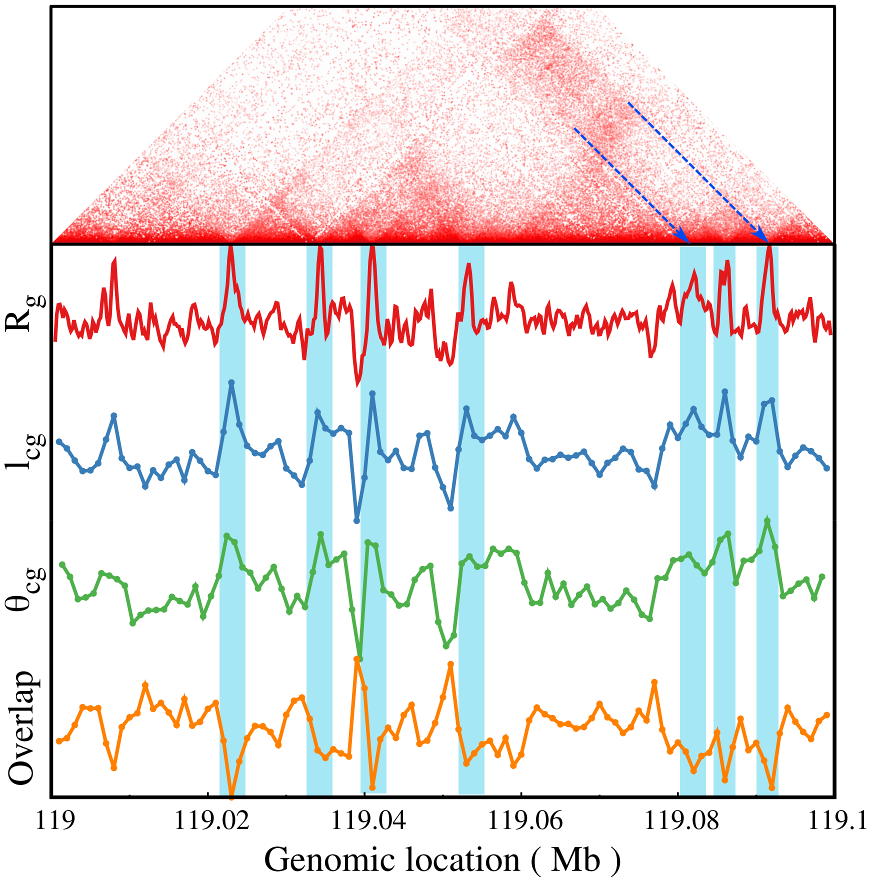


FIG. S6 The $R_g$, $l_{cg}$, $\theta_{cg}$, and overlap parameter is plotted as a function of genomic location for Cbx8 locus. Here we observe different polymer properties at the boundary of heterochromatic domains.

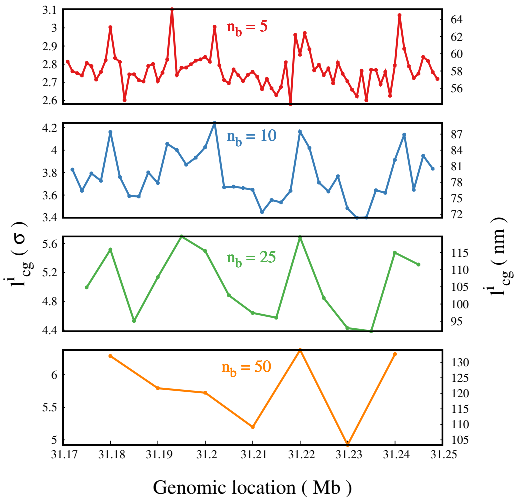


(a) Ppm1g locus

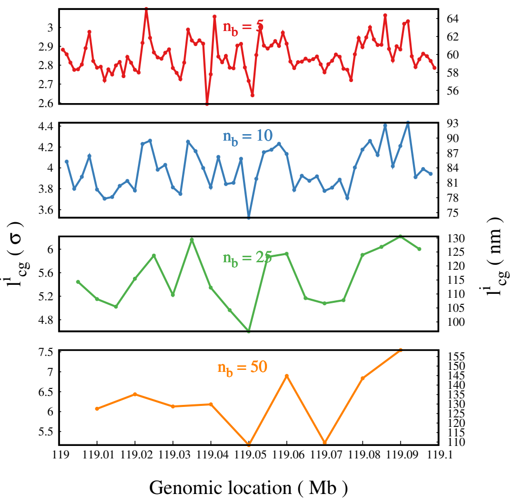


(b) Cbx8 locus

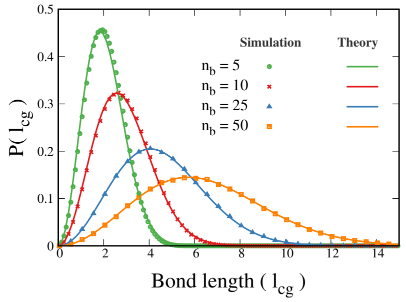


(c) Ideal chain

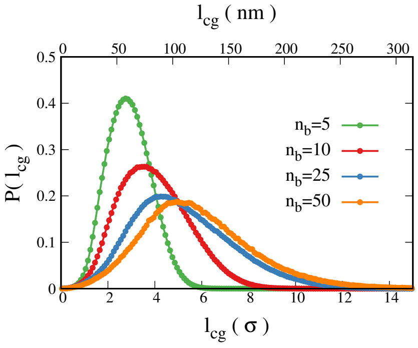


(d) Gm29683 locus

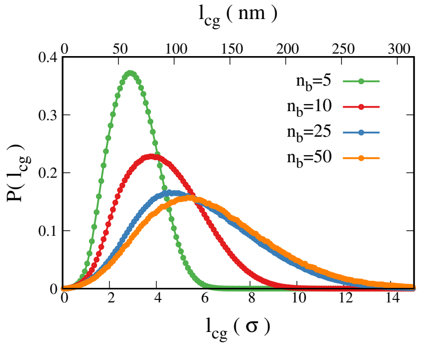


(e) Arsg locus

FIG. S7 Average bond length ($l_cg$) between neighboring CG beads as a function of genomic location for (a) Gm29683 locus, (b) SAW and (c) globular polymer. Distribution of $R_g$ for (d) Arsg locus and (e) Gm29683 locus.

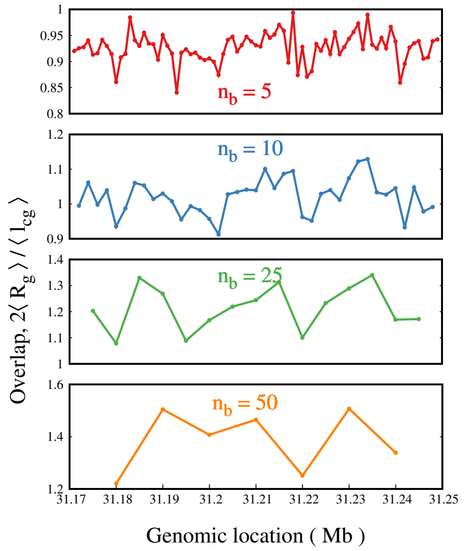


(a) Ppm1g locus

(b) Gm29683 locus

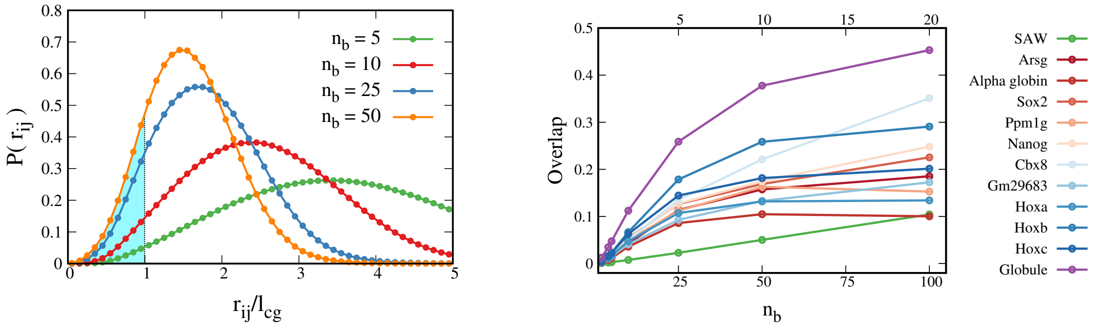


(c)

(d) Overlap between non-bonded beads

FIG. S8 (a) Distribution of bond length for ideal chain simulation fitted with the analytical expression form Laso et al. (1991). Our simulation results compare well with the analytical formula validating our simulation methods. Distribution of bond length for different values of $n_b$ is plotted for (b) Gm29683 locus and (c) Arsg locus. Overlap is plotted as function of genomic location for different values of $n_b$ for (d) the Ppm1g locus and (e) Gm29683 locus. (f) The distribution of 3D distance between non-bonded ($|i - j| > 1$) coarse-grained beads is plotted for different coarse-graining levels (Arsg locus). The area under the curve till $r_{ij} < l_{cg}$ (shaded region) can be used as a measure for the overlap or softness of the coarse-grained beads, which is plotted as a function of $n_b$ for various regions in (g).

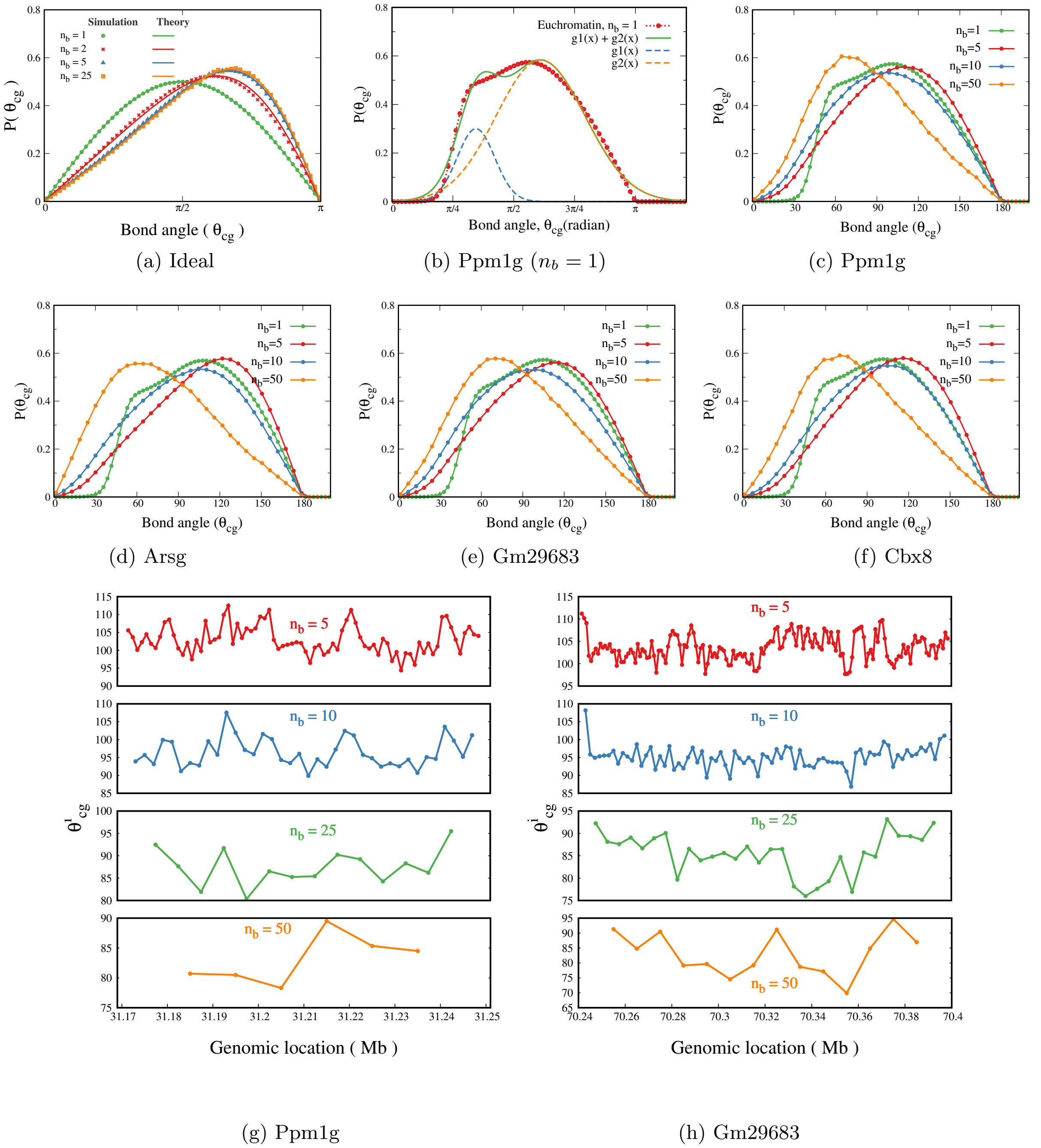


(a) Ideal

(b) Ppm1g ($n_b = 1$)

(c) Ppm1g

(d) Arsg

(e) Gm29683

(f) Cbx8

(g) Ppm1g

(h) Gm29683

FIG. S9 (a) Distribution of bond angle for ideal chain simulation fitted with the analytical expression form Laso et al. (1991). (b) Bond angle distribution from simulation for Ppm1g locus ($n_b = 1$) is fitted with two gaussian curves with mean 62° and 109.7°. Here $g1(x) = 0.3\ e^{-\left((x-1.083)/0.34\right)^2}$ and $g2(x) = 0.58\ e^{-\left((x-1.92)/0.81)\right)^2}$. Angle distribution for various values of $n_b$ is plotted for (c) Ppm1g, (d) Arsg, (e) Gm29683, and (f) Cbx8 loci. Average bond angle between CG beads as a function of genomic location for (g) Ppm1g locus and (h) Gm29683 locus.

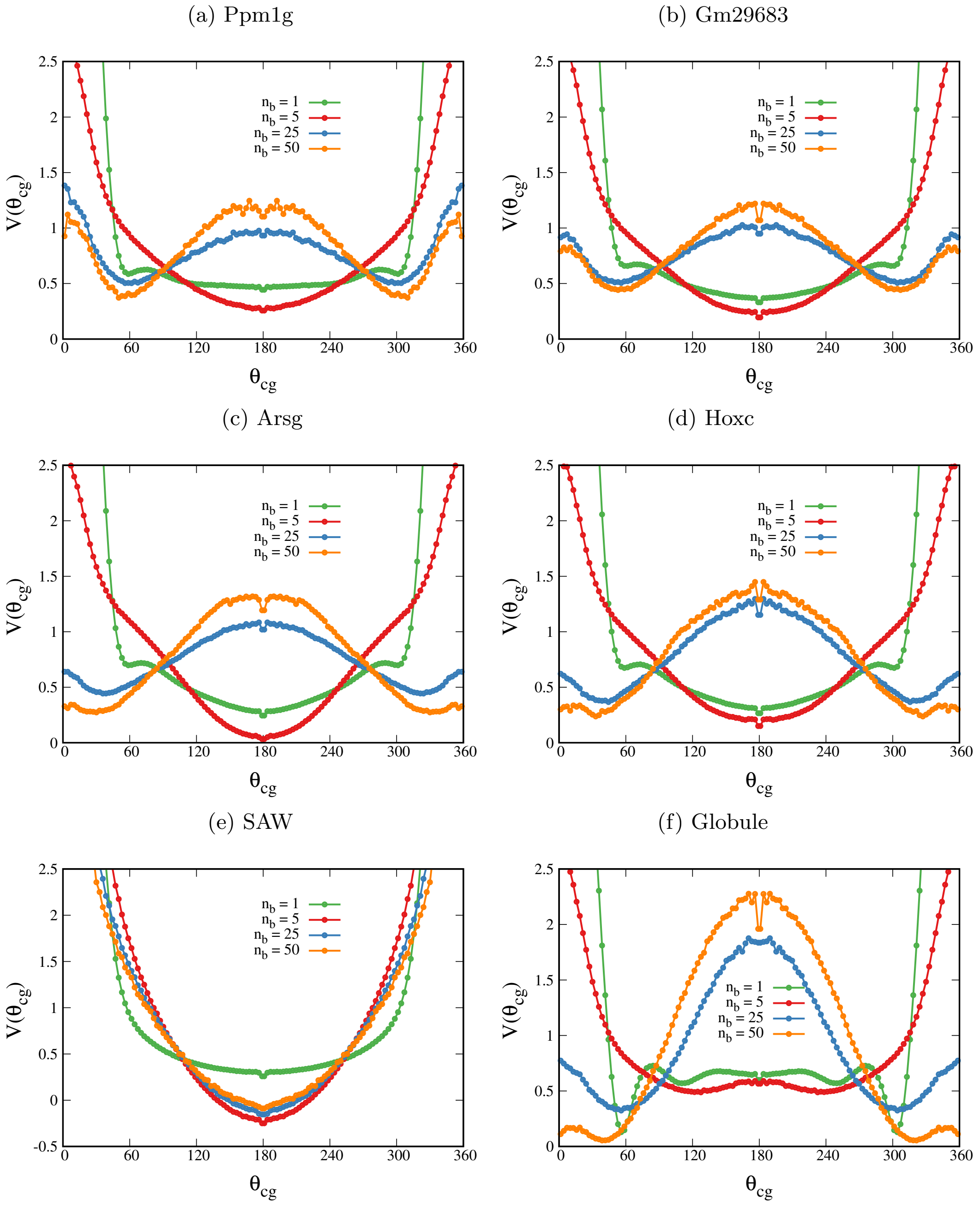


FIG. S10 Bond angle potential for different levels of coarse-graining is plotted for (a) Ppm1g locus, (b) Gm29683 locus, (c) Arsg locus,(d) Hoxc locus,(e) SAW polymer, and (f) globule polymer. $\theta_{cg} = 180^\circ$ represents straight configuraiont while $\theta_{cg} = 0^\circ, 360^\circ$ represents completely bent configuration. Note that the angle potential from $0^\circ$ to $180^\circ$ is reflected to $180^\circ$ to $360^\circ$ for representation purposes as potential is symmetric around $180^\circ$

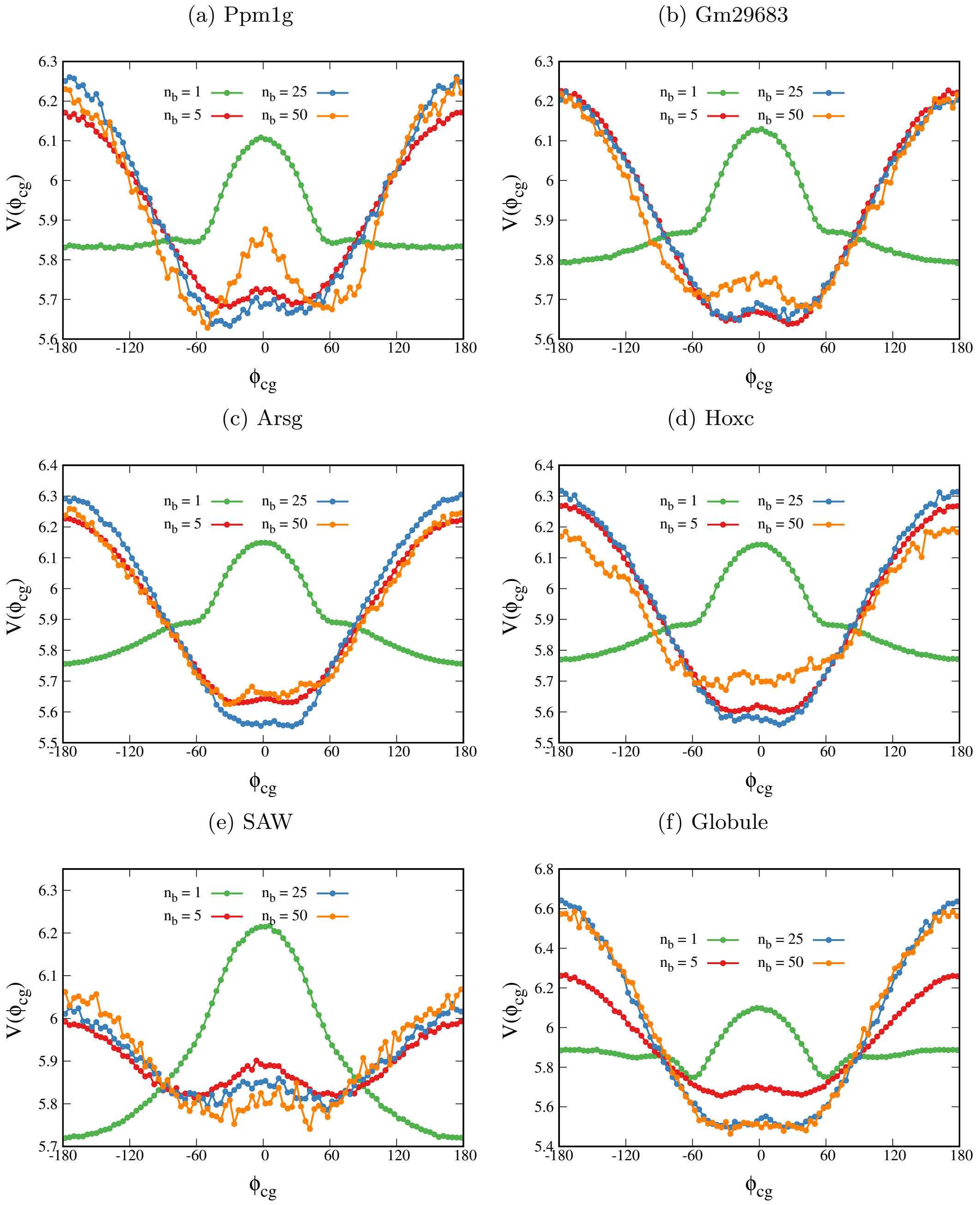


FIG. S11 Dihedral angle potential for different levels of coarse-graining is plotted for (a) Ppm1g locus, (b) Gm29683 locus, (c) Arsg locus,(d) Hoxc locus,(e) SAW polymer, and (f) globule polymer.

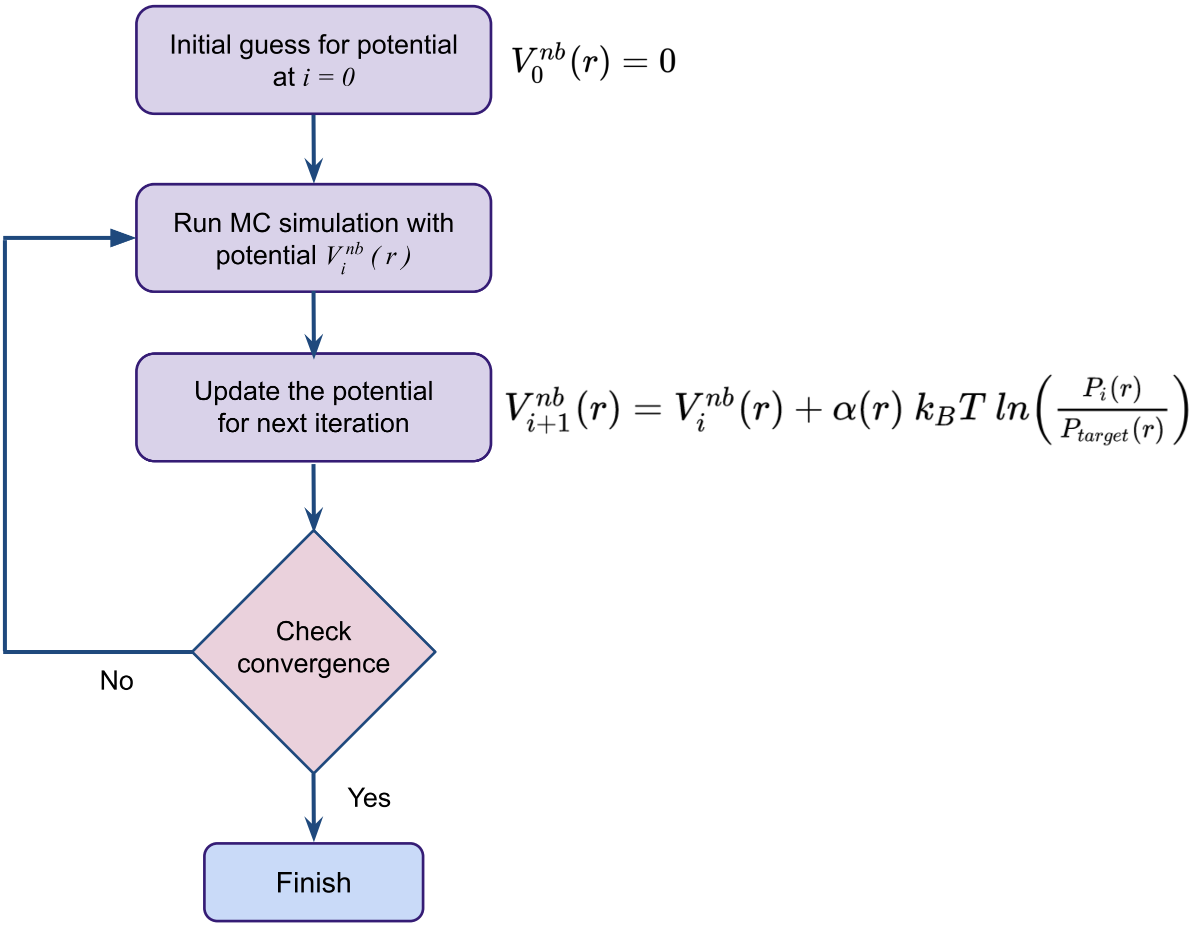


FIG. S12 Flow-chart explaining the iterative Boltzmann inversion algorithm.

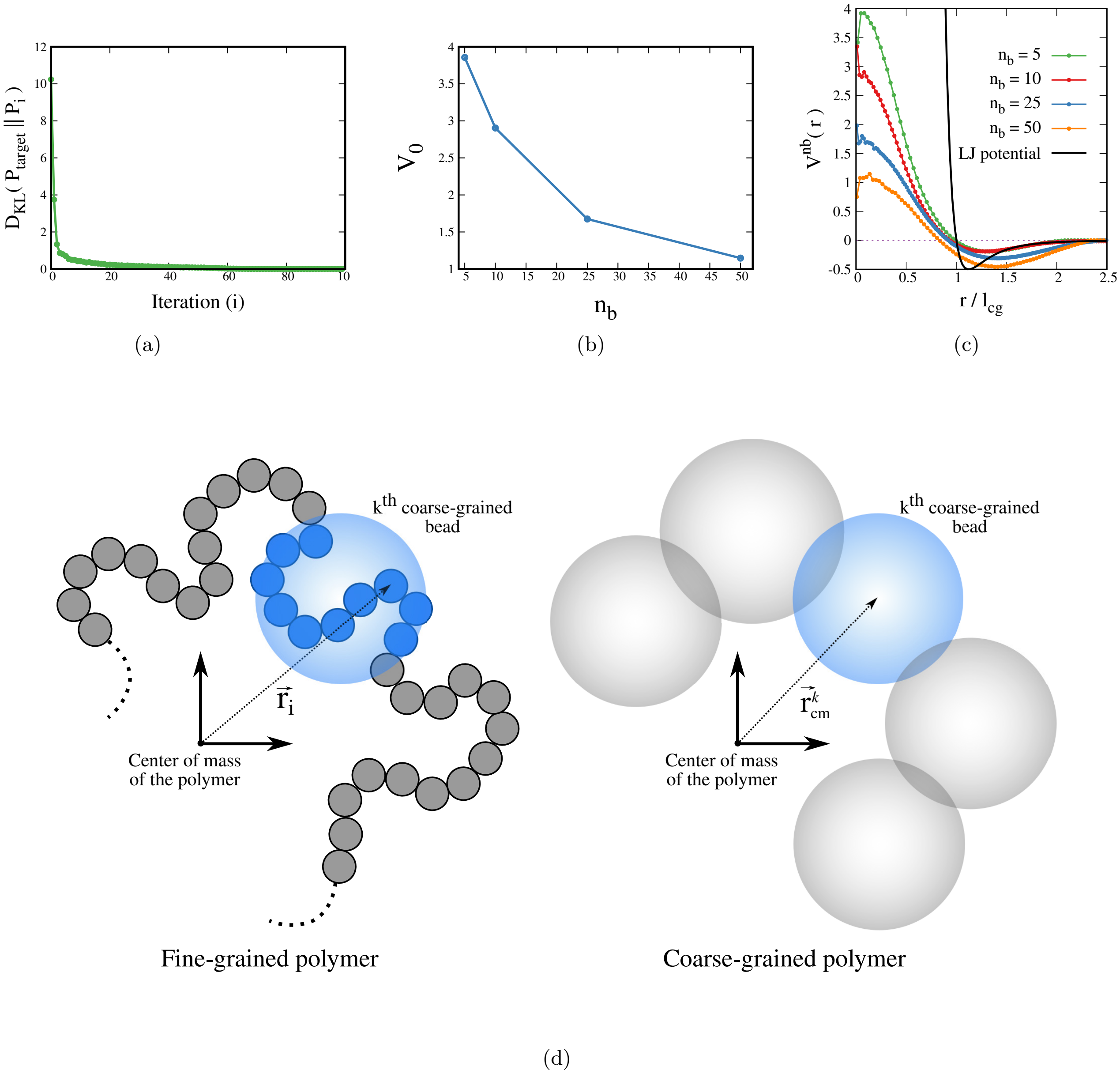


FIG. S13 (a) The Kullback-Leibler Divergence between the target distribution and the distribution from CG model is plotted as function of iteration $i$ for $n_b = 50$. (b) The height of the potential $V^{nb}$ around $r = 0$ is plotted as a function of coarse-graining. (c) Comparison of the soft potential obtained from the iterative Boltzmann inversion method with Lennard-Jones potential ($\epsilon = 0.5, \sigma = l_{cg}$). (d) The schematic showing the fine-grained polymer and coarse-grained polymer in center of mass frame of reference.

**The radius of gyration of a polymer decreases slightly with increasing coarse-graining:**

The radius of gyration of a polymer is a measure of its size and is defined as the root mean square distance of the beads from their center of mass. When we coarse-grain a polymer, we effectively replace a group of $n_b$ beads with a single effective bead located at the center of mass of the group. The distance between the effective bead and the center of mass of the polymer is smaller than the root mean square distance between the original fine-grained beads and the center of mass of the polymer (see below). As a result, the radius of gyration decreases slightly as we coarse-grain the polymer.

To illustrate this, consider the fine-grained polymer consisting of $N$ beads in the polymer's center of mass frame of reference (see Fig. S13(d)). The corresponding coarse-grained polymer consists of $N/n_b$ beads, with each CG bead replaced by the center of mass of $n_b$ fine-grained beads. Let us consider the contribution of the blue segment of fine-grained beads and the corresponding CG bead in the radius of gyration calculation. The root mean square distance between fine-grained beads in $k^{th}$ segment and the center of mass of the polymer is given by

$$R_{FG} = \sqrt{\frac{1}{n_b} \sum_{i=n_b k}^{n_b k + n_b} r_i^2} \quad .$$

Here, $r_i$ is the distance of $i^{th}$ bead from the center of mass of the polymer. The corresponding distance of the CG bead will be the distance of the blue segment's center of mass from the polymer's center of mass (i.e., origin). This can be written as

$$R_{CG} = r_{cm}^k = \frac{1}{n_b} \sum_{i=n_b k}^{n_b k + n_b} r_i \quad .$$

Since the root-mean-square of any quantity is always greater than or equal to the mean, mathematically, $R_{CG}$ is always less than $R_{FG}$. This applies to all other CG beads. Hence, the radius of gyration of the CG polymer will be slightly smaller than that of the fine-grained polymer.
